# Transgenic αβ TCR tonic signaling is leukemogenic while strong stimulation is leukemia-suppressive

**DOI:** 10.1101/2024.01.17.576006

**Authors:** Telmo A. Catarino, Ivette Pacheco-Leyva, João L. Pereira, Marina Baessa, Nuno R. dos Santos

## Abstract

The pre-T cell receptor (TCR) and TCR complexes are frequently expressed in T-cell acute lymphoblastic leukemia (T-ALL), an aggressive T cell precursor malignancy. Although mutations in TCR components are infrequent in T-ALL, earlier research indicated that transgenic αβ TCR expression in mouse T cell precursors promoted T-ALL development. However, we recently found that stimulation of TCR signaling in T-ALL induced leukemic cell apoptosis and suppressed leukemia. Our aim was to elucidate if a given αβ TCR complex has a dual role in leukemogenesis depending on the nature of the stimulus. We demonstrate that transgenic expression of the Marilyn αβ TCR, specific for the H-Y male antigen presented by major histocompatibility complex class II, triggers T-ALL development exclusively in female mice. This T-ALL exhibited *Notch1* mutations, *Cdkn2a* copy number loss, immature immunophenotype and infiltrated both lymphoid and non-lymphoid organs. Furthermore, leukemic cells expressed surface CD5, a marker of tonic TCR signaling. T-ALL efficiently developed in *Rag2*-deficient Marilyn transgenic females, indicating that Rag2-mediated recombination is not implicated in this T-ALL model. Remarkably, exposure of Marilyn female T-ALL to male antigen in recipient mice resulted in T-ALL apoptosis and prolonged mouse survival. These findings underscore that the same αβ TCR complex has a dual role in T-ALL in that its tonic stimulation is leukemogenic, while strong stimulation is leukemia-suppressive.

## Introduction

The pre-T cell receptor (TCR) and αβ TCR are critical protein complexes for T cell development. Pre-TCR expression in CD4^−^CD8^−^ double-negative (DN) thymocytes induces their survival, proliferation and maturation through the β-selection checkpoint,^1,2^ whereas surface expression of the αβ TCR in CD4^+^CD8^+^ double-positive (DP), thymocytes drives thymic positive and negative selection to establish major histocompatibility complex (MHC) restriction and eliminate autoreactive thymocytes. ^3^ The pre-TCR, TCR and CD3 proteins are frequently expressed in T-cell acute lymphoblastic leukemia (T-ALL). ^4^ Although T-ALL is genetically a very heterogeneous disease, with mutations detected in multiple genes, a number of genetic alterations have been recurrently reported for T-ALL, including *CDKN2A* loss of function, *NOTCH1*-activating mutations, and mutations in JAK/STAT signaling pathway genes (e.g. *IL7R*, *JAK3*, *JAK1*, and *STAT5B*).^5^ Mutations in components of the TCR signaling machinery are rare, ^5^ but the Lck kinase, a key member of the pre-TCR and TCR signaling pathways, has been shown to be activated in T-ALL and put forward as a potential therapeutic target. ^6–9^ These observations hint that pre-TCR signaling can promote human T cell leukemogenesis. In this line, genetic inactivation of the pre-TCR complex (e.g. CD3 ε, pT α or Rag1/2 deficiency) was shown to delay T-ALL development in several mouse models.^10–14^ In contrast, the impact and function of TCR signaling in T-ALL development and progression remains controversial. While transgenic TCR expression was shown to promote mouse T-ALL development in cooperation with STAT5 overexpression or deficiency of p53 or Tpl2 proteins, ^15–17^ expression of endogenous αβ TCR or MHC-dependent antigen presentation were not required for leukemogenesis in the ETV6::JAK2 or TAL/LMO transgenic mouse models. ^13,14^ Indicating that TCR could rather have an antagonistic action in T-ALL, T-ALL onset in a *Pten*-deficient mouse model was accelerated by TCR genetic inactivation and delayed by expression of transgenic TCR. ^18^ In this line, it was also reported that strong agonist TCR stimulation of mouse ETV6::JAK2-driven T-ALL or patient T-ALL was anti-leukemic and induced a transcriptional program similar to that involved in thymocyte negative selection. ^19^ Overall, these observations suggest that, depending on the oncogenic drivers of leukemia and degree of TCR stimulation, the TCR signaling complex might play either a leukemia-promoting or a leukemia-suppressive role.

Here, our aim was to investigate the role of basal versus strong TCR signaling in T-ALL leukemogenesis. In normal T cells, TCR recognition of self-peptides presented by MHC (self-pMHC) induces tonic signaling, while agonist peptides induce strong TCR signaling. Using the Marilyn transgenic TCR mouse strain, specific for the H-Y male antigen, we show that tonic TCR signaling led to *Notch1*-mutated T-ALL exclusively in females. MHC-dependent agonist antigenic stimulation prevented spontaneous T-ALL development in males, induced apoptosis of female leukemic cells and hampered their leukemogenic potential. Our results support the notion that αβ TCR tonic signaling promotes T cell precursor malignant transformation and that strong signaling suppresses leukemia.

## Results

### Transgenic expression of an αβ TCR specific for the H-Y male antigen drives leukemia development in female mice

To study the effect of TCR expression in leukemia development, we resorted to Marilyn transgenic mice, which express an MHC class II-restricted transgenic TCR (TCR-Vα 1.1/Vβ 6) specific for the H-Y male antigen (HY-TCR henceforth). ^20^ In this model, male thymocytes expressing the HY-TCR are exposed to the cognate antigen (DBY peptide), through MHC class II presentation, and are eliminated by apoptosis. In contrast, HY-TCR-expressing female thymocytes undergo positive selection and differentiate as mature CD4^+^ T cells. To verify if the transgenic Marilyn HY-TCR promoted leukemogenesis, we bred transgenic mice expressing the ETV6::JAK2 fusion protein in lymphoid cells with Marilyn mice. As expected, ^13^ all ETV6::JAK2 transgenic mice developed T-ALL expressing CD8, CD25, CD24, CD3ε and TCRβ (not shown). Remarkably, co-expression of the Marilyn HY-TCR with ETV6::JAK2 accelerated spontaneous T-ALL development in females (median survival of 10.5 weeks versus 14 weeks; Figure 1A). This suggests that HY-TCR tonic signaling, which is induced by TCR interaction with self-peptides presented by MHC, cooperates with the ETV6::JAK2 transgene in driving leukemogenesis.

**Figure 1.**
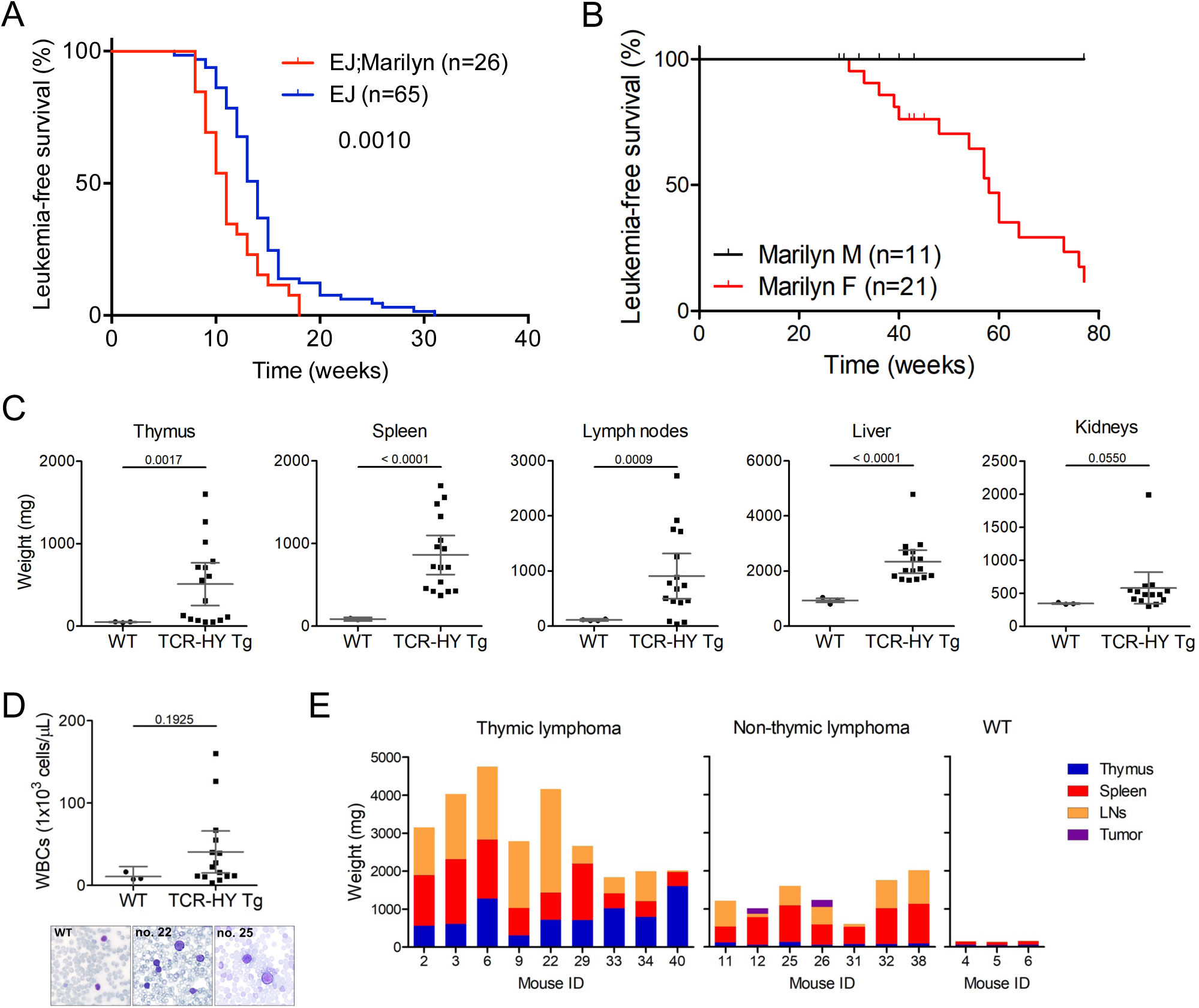
HY-TCR transgene expression drives leukemia development. (A,B) Kaplan-Meier leukemia-free survival curves of ETV6::JAK2 (male and female together) and female ETV6::JAK2;Marilyn transgenic mice (A) and male (M) and female (F) Marilyn transgenic mice (B). Log-rank test *P* values are shown in graphs. (C) Organ weights of Marilyn females with leukemia/lymphoma compared to those of 7 month-old wild-type (WT) females (n=3), or 10-week-old WT males for lymph nodes (n=4). For Marilyn females n=16, except n=14 for kidneys. (D) White blood cell (WBC) counts of Marilyn (n=15) and WT (n=3) females. Lower panels show blood smears of representative WT and diseased Marilyn females. *P* values in (C) were obtained from unpaired *t* tests with Welch’s correction, and in (D) from Mann-Whitney test. Mean and standard error of the mean (SEM) are shown. (E) Histogram representation of tumor burden in lymphoid organs (thymus, spleen, and lymph nodes (LNs)) of Marilyn females with or without thymic lymphoma. Female no. 12 had a mediastinal lymphoid tumor, while no. 26 had lymphoid tumors on the duodenum wall. Each column represents a mouse. Wild-type (WT) female mice were used as control.

To determine whether HY-TCR tonic signaling alone could induce T cell malignant transformation, we monitored a cohort of both Marilyn females and males until 18 months of age. Remarkably, while most females developed fatal hematological disease (median survival of 58 weeks), none of the males did (Figure 1B). Macroscopic analyses showed that Marilyn females developed enlarged lymphoid (thymus, spleen and lymph nodes (LNs)) and non-lymphoid organs (mainly liver and kidneys) (Figure 1C; Supplementary Figure 1), together with high number of blood lymphoblasts (Figure 1D). Histological analyses revealed massive lymphoid cell infiltration of liver, kidneys, lungs and bone marrow (Supplementary Figure 2). These data indicate that tonic (in females) but not strong (in males) transgenic HY-TCR signaling is sufficient to induce leukemia and lymphoma development.

### Transgenic HY-TCR expression induced T-cell disease with variable immunophenotype

Macroscopic analysis revealed two distinct groups of diseased Marilyn females: a subset with thymic lymphoma (TL; 9 cases) and a subset characterized by splenomegaly and lymphadenopathy without thymic lymphoma (non-TL; 7 cases; Figure 1E). TL females developed leukemia slightly faster and presented higher total lymphoid organ disease burden than non-TL females, suggestive of a more aggressive disease course (Supplementary Figures 3A,B). For a better characterization of the hematological disease developing in different Marilyn females, we performed immunophenotyping by flow cytometry. To define the T cell maturation status, we probed expression of CD24, which is downregulated upon thymocyte maturation, CD5, which is upregulated after thymocyte positive selection, and CD44, which is expressed in memory T cells. ^21,22^ Within the TL subgroup, five mice (nos. 2, 6, 22, 33 and 40) presented thymic T cells with immature phenotype, characterized by surface expression of CD90.2, TCR-V β 6, and CD5, low levels of CD4 and CD8, high CD24 levels, and CD44 negativity (Figure 2 and Supplementary Table 1). Such immunophenotype was recapitulated in leukemic cells disseminated to the spleen, LNs, and bone marrow (Supplementary Figure 4 and not shown). Two other TL cases (nos. 9 and 29) presented a more mature T cell immunophenotype, characterized by CD4^+^CD8^−^TCR-Vβ6^+^CD5^+^ leukemic cells with CD44 expression (Figure 2). These cells were also present in spleen and lymph nodes (Supplementary Figure 4). Finally, and rather unexpectedly, two TL cases (nos. 3 and 34) presented a large proportion of B220-positive B cells, concomitantly with TCR-V β 6^+^CD5^+^CD44^+^ CD4 single-positive (SP) and CD8SP T cells (Figure 2). Of note, TL cases with immature immunophenotype, but not those with mature phenotype, could reinitiate disease in recipient syngeneic mice (Supplementary Table 1), suggesting that only the former were malignantly transformed.

**Figure 2.**
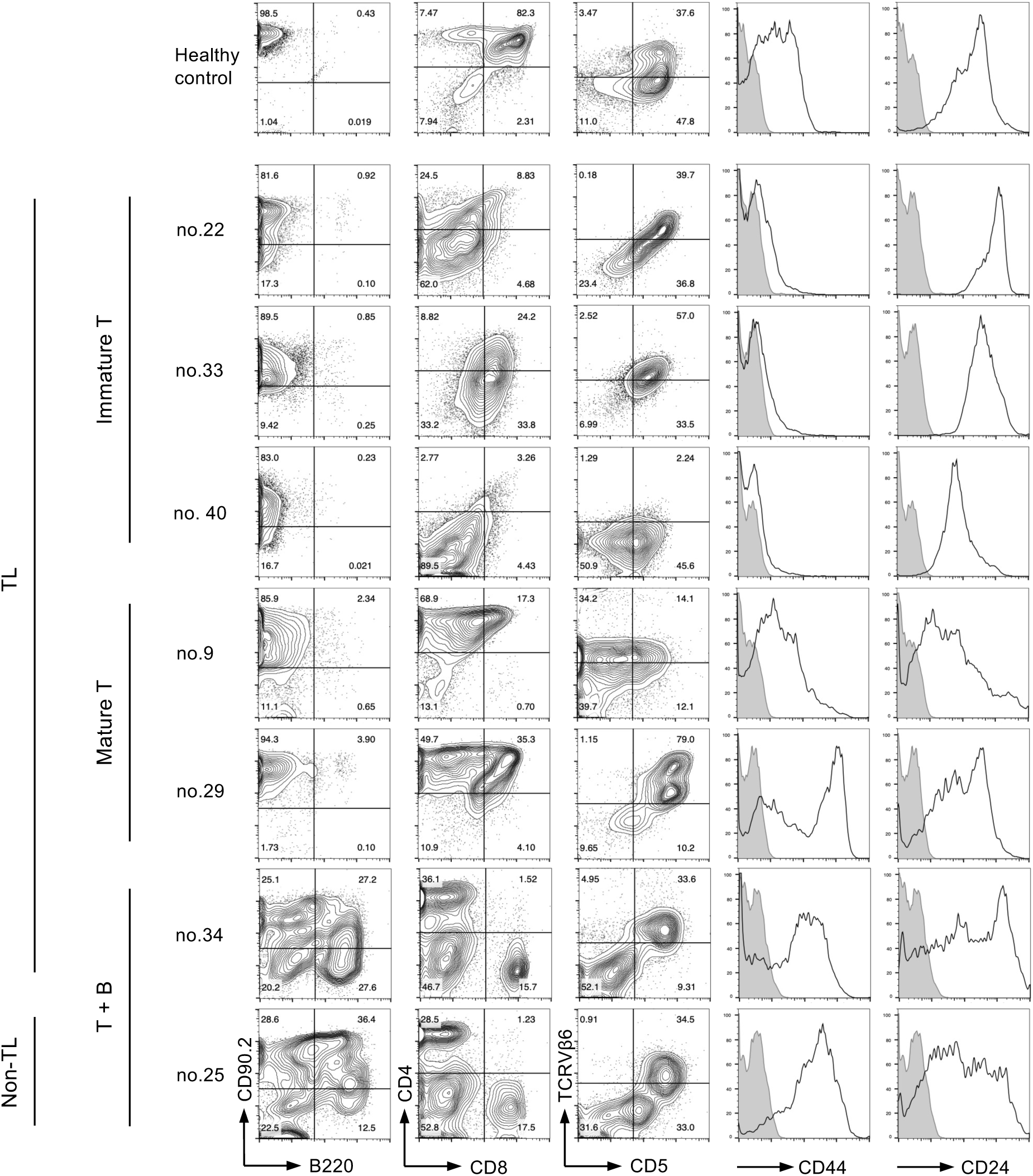
Immunophenotypic analysis of HY-TCR leukemic cells. Flow cytometry analysis of thymus-derived leukemic cells from representative Marilyn mice presenting immature T cell phenotype, based on CD44 negativity and high CD24 expression (nos. 22, 33 and 40), mature T cell phenotype, based on CD44 positivity and CD24 intermediate/low expression (nos. 9 and 29), and mixed B and T cell phenotype, based on B220 and CD90.2 expression (nos. 25 and 34). In CD44 and CD24 histogram plots, unstained cells were used as negative control (grey shades). Healthy HY-TCR female total thymocyte immunostaining is shown as control. TL, thymic lymphoma.

Non-TL Marilyn females exhibited a mixture of B220^+^ B cells and CD90.2^+^TCR-Vβ6^+^CD5^+^CD44^+^ CD4SP cells in lymphoid organs (Figure 2 and Supplementary Figure 4 and Table 1). Analysis of CD69 expression, a marker of TCR stimulation, ^21^ showed that TCR-V β 6^+^ cells from mixed B and T lymphomas were CD69-positive, indicating they experienced recent TCR stimulation (Supplementary Figure 5). This suggests that mixed B and T disease found in Marilyn female mice was driven by continuous stimulation of the HY-TCR transgene expressed in mature CD4SP or CD8SP T cells. Immature-type leukemic cells did not express CD69, suggesting that their proliferation was likely driven by oncogenic mechanisms rather than TCR stimulation.

We conclude that while one third (5 of 16) of Marilyn TCR transgenic females developed a transplantable immature disease, akin to T-ALL, marked by pronounced thymic enlargement, the remaining diseased mice developed a non-transplantable and heterogeneous mature T and B nonmalignant lymphoproliferation affecting the thymus, spleen, lymph nodes and other organs, with statistically significant longer delay (median survival of 40 versus 60 weeks; Supplementary Figure 6).

### Marilyn T-ALL presents *Notch1* PEST domain mutations and genetic copy number alterations

To determine whether Marilyn T-ALL was associated with secondary genetic alterations, we first sequenced the *Notch1* exon 34 (PEST domain), where most murine T-ALL *Notch1* mutations have been found. ^23,24^ Four out of 5 Marilyn T-ALL cases analyzed carried exon 34 frameshift mutations (Supplementary Figure 7 and Table 1), while none were detected in cases of mixed T and B cell disease (Supplementary Table 1). Additionally, by performing low-coverage whole-genome sequencing, we found that T-ALL samples had near-diploid genomes, with only a few copy number alterations: a chromosome 15 trisomy (case no. 6), a chromosome 10 trisomy (case no. 33) (Supplementary Figure 8A), and loss of the *Cdkn2a* locus (chromosome 4) in three cases (nos. 2, 22 and 40) (Supplementary Figure 8B). In contrast, no consistent copy number alterations were found in 2 cases of mixed T and B cell disease (Supplementary Figure 8A). Deletions of the *Tcra/d*, *Tcrb* and *Tcrg* loci were also found in Marilyn T-ALL (Supplementary Figure 8C), indicating that these loci underwent genetic rearrangements and that leukemia arose from post-β-selection thymocytes with rearranged TCR loci. As expected from their T lineage origin, T-ALL cases did not exhibit consistent copy number alterations in the immunoglobulin *Ighm* or *Igk* loci (Supplementary Figure 8C). The detection of clonal copy number genetic alterations in Marilyn T-ALL bulk populations further underscores their malignant nature. Given the high frequency of *NOTCH1*-activating mutations and *CDKN2A* inactivation in human T-ALL, ^5^ our findings show that the Marilyn transgenic HY-TCR T-ALL recapitulates key cellular and molecular features of the human disease.

### HY-TCR-induced T-ALL is not dependent on *Rag2* recombinase expression

Since several diseased Marilyn females developed B cell lymphoproliferation, we generated Marilyn mice with *Rag2* constitutive knockout to block B cell development, while maintaining HY-TCR-dependent T-cell development. This approach simultaneously assessed whether Rag-mediated gene recombination plays a role in HY-TCR-induced T cell leukemogenesis. Marilyn HY-TCR cohorts with (Marilyn;*Rag2*^+/−^) or without Rag2 protein (Marilyn;*Rag2*^−/−^) were generated in another animal facility (i3S/Porto). Surprisingly, most Marilyn;*Rag2*^+/−^ female mice (78%) did not develop disease, with the exception of two mice with splenomegaly at nearly 18 months of age, which was composed of B and myeloid cells (Figure 3A and not shown). Although half of Marilyn;*Rag2*^−/−^ males developed hematological splenic disease after 1 year of age (4 cases of myeloid proliferation and 1 case of *Notch1*-mutated TCR-V β 6^+^ CD4^−^CD8^−^ splenic lymphoma), nearly all Marilyn;*Rag2*^−/−^ females developed lymphoma with much earlier onset than males (median survival of 46 and 70 weeks, respectively; Figure 3A). All diseased Marilyn;*Rag2*^−/−^ females presented thymic lymphoma, of variable size, frequently accompanied by enlarged spleen, LNs, liver and kidneys (Figure 3B; Supplementary Figure 9A). Leukemias from Marilyn;*Rag2*^−/−^ females were mostly composed of CD4^+/low^CD8^+/low^ immature T cells expressing TCR-V β 6, high levels of CD24, CD44 negativity and variable expression of CD25 and CD5 (Figure 3C, panel i, and Supplementary Table 2). A similar immunophenotype was detected across different affected organs, suggesting that diseased cells disseminated from the thymus to other organs (Figure 3C, panel ii). The pathological and immunophenotypic features, together with the observed 60% frequency of *Notch1* exon 34 frameshift mutations and ability to reinitiate disease in syngeneic recipients (Supplementary Table 2), indicate that Marilyn;*Rag2*^−/−^ females developed T-ALL. To assess the degree of leukemic cell infiltration in different Marilyn;*Rag2*^−/−^ organs, we performed histological analysis. Lymphoblasts were conspicuous in the thymus and spleen, and infiltrated with variable extent, the liver, kidneys, lungs and bone marrow (Supplementary Figure 9B). Interestingly, immunohistochemical analyses of thymus histological sections showed that while Marilyn T-ALL cells were highly proliferative, with conspicuous Ki67 staining, they maintained a high level of apoptosis, as gauged by cleaved caspase 3 staining (Figure 3D). In sum, these data confirm that Marilyn HY-TCR tonic signaling drives T-ALL and demonstrated that Rag recombinase-mediated DNA rearrangements are not implicated.

**Figure 3.**
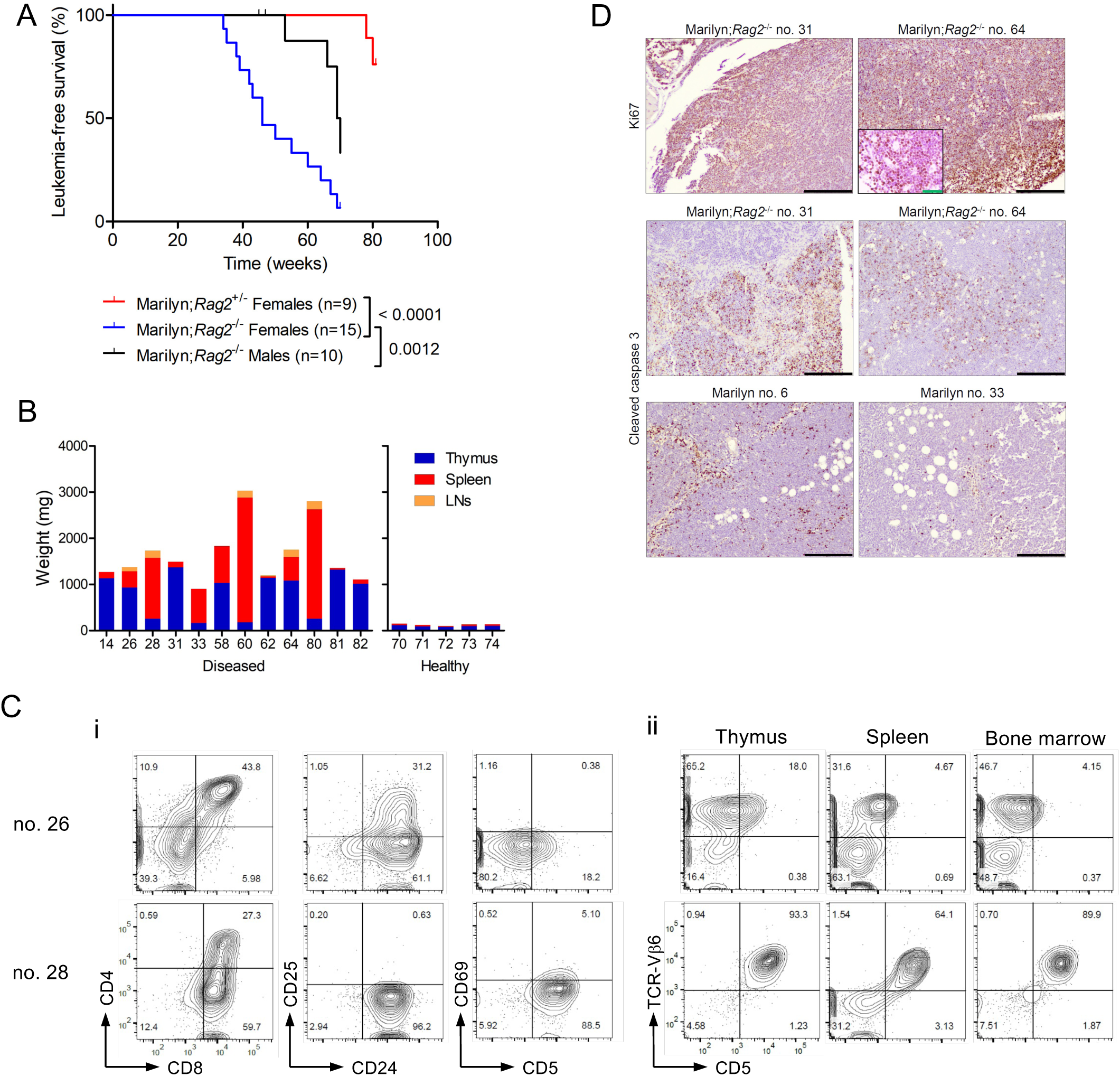
Transgenic HY-TCR expression in *Rag2^−/−^*mice drives development of T-ALL with thymic lymphomas. (A) Kaplan-Meier leukemia-free survival curve for the indicated genotypes and number of mice (n). Log-rank test *P* values are indicated. (B) Histogram representation of lymphoid organ weight (thymus, spleen and lymph nodes (LNs)) of Marilyn females with leukemia (left graph) and healthy (right graph). Each column represents a mouse. (C) Flow cytometry immunostaining of cells from the thymus (i) and indicated organs (ii) of two representative Marilyn;*Rag2^−/−^*leukemic mice with the indicated surface markers. (D) Immunohistochemical Ki67 and cleaved-caspase 3 staining of representative thymic lymphoma sections from Marilyn;*Rag2^−/−^* and Marilyn;*Rag2^+/+^* diseased mice. Black scale bars: 500 μm; green scale bar: 200 μm.

### Hematological disease in HY-TCR transgenic females is associated with microbial status

Marilyn females in the UAlg/Faro animal facility had higher incidence of hematological disease than the Marilyn;*Rag2*^+/−^ females in the i3S/Porto facility. Since opportunistic microorganisms were detected in the UAlg but not the i3S facility (see Materials and methods), we posit that an environmental input present only at the UAlg facility increased susceptibility for lymphoid disease. Considering that bacteria are frequently present in tumors, and can be detected by immunohistochemical methods, ^25^ we probed lymphomas obtained in both facilities for the presence of bacterial-derived lipopolysaccharide (LPS). Indeed, Marilyn lymphomas derived from the UAlg facility displayed more conspicuous LPS immunostaining than Marilyn;*Rag2*^−/−^ lymphomas derived from the i3S facility (Supplementary Figure 10A). Of note, the pattern of LPS staining in Marilyn splenic lymphomas overlapped with B220^+^ B cell areas (Supplementary Figure 10B). These data suggest that higher bacterial exposure could have promoted hematological diseases in Marilyn females.

### The transgenic HY-TCR is functional in Marilyn T-ALL cells and its stimulation delays leukemogenesis

Given that antigenic stimulation prevented spontaneous T-ALL development in male HY-TCR mice, we set out to determine the response of female HY-TCR leukemic cells to TCR stimulation. Marilyn T-ALL cells readily increased CD69 and CD5 expression and enlarged in size when stimulated *in vitro* with plate-bound CD3 antibody, which directly activates the TCR signaling complex, or phorbol 12-myristate-13-acetate (PMA) and ionomycin, triggering downstream activation of protein kinase C and intracellular calcium pathways (Supplementary Figure 11). Previous research demonstrated that agonist TCR stimulation can induce T-ALL apoptosis *in vitro*. ^19^ However, Marilyn leukemic cells underwent extensive apoptosis *in vitro*, impeding determination if TCR stimulation induced cell death. To assess the impact of TCR stimulation *in vivo*, Marilyn T-ALL cells were injected in syngeneic male and female mice and disease progression followed by peripheral blood collection. Marilyn T-ALL cells were found in the blood of females, but not males, as early as 30 days post-injection (Figure 4A). Most female recipients rapidly developed fatal T-ALL, while male recipients often failed to develop leukemia and therefore survived longer (Figure 4B). In addition, females infused with Marilyn T-ALL presented significantly larger spleen and kidneys (Figure 4C), and more extensive BM leukemia infiltration (Figure 4D), than male recipients euthanized simultaneously. Furthermore, male spleens showed higher levels of cleaved caspase 3 immunostaining than female spleens (Figure 4E), indicating that delayed leukemogenesis in males was caused by apoptosis induction.

**Figure 4.**
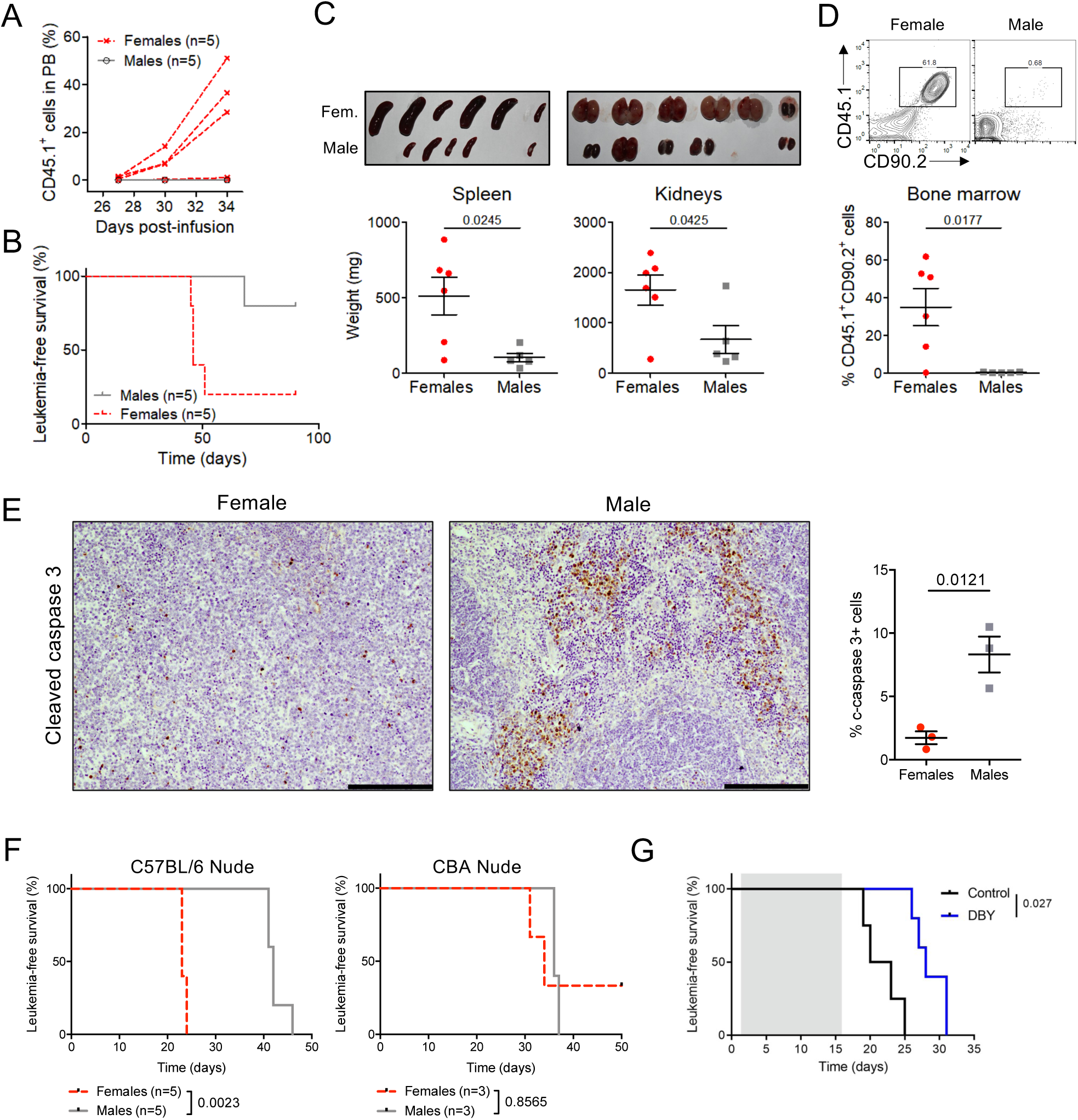
HY-TCR T-ALL antigenic stimulation *in vivo* hampers leukemia development. (A) Percentage of CD45.1^+^ T-ALL cells detected by flow cytometry in the peripheral blood (PB) of male and female C57BL/6 mice infused with female HY-TCR T-ALL cells. *P* value was obtained with two-way ANOVA. (B) Kaplan-Meier leukemia-free survival curve for the same mice as in (A), which is representative of two experiments performed with different primary leukemias. *P* value obtained with log-rank test. (C) Spleens and kidneys (top panels) and respective organ weights (bottom panels) for male (n=5) and female (n=6) C57BL/6 mice 21 days after infusion with HY-TCR T-ALL cells. (D) Percentage of CD45.1^+^CD90.2^+^ HY-TCR T-ALL cells present in the bone marrow of males and females shown in (C). *P* values in (C) and (D) were obtained from unpaired *t* tests with Welch’s correction. (E) Representative sections and quantification of anti-cleaved caspase 3 immunohistochemistry for spleens from females and males shown in (C,D; n=3). *P* value from unpaired *t* test. (C-E) Graphs show mean and SEM. (F) Kaplan-Meier leukemia-free survival curves for syngeneic C57BL/6 nude or allogeneic CBA nude male and female mice injected with female Marilyn T-ALL cells. Number of mice and *P* values from log-rank test are indicated. (G) Kaplan-Meier leukemia-free survival curves for syngeneic *Rag2*^−/−^ females infused on day 0 with female Marilyn T-ALL cells and treated once a week with DBY peptide (0.01 mg; n=5) or Control (n=4). Gray shade represents treatment duration of 2 weeks.

To address whether the leukemia-delaying effect of agonist-induced TCR stimulation in Marilyn T-ALL cells was dependent on MHC class II presentation, Marilyn T-ALL cells were infused in immunodeficient syngeneic and allogenic males and females. In contrast to the delayed T-ALL development in syngeneic males, as compared to female counterparts (Figure 4F, left panel), no significant difference between allogeneic males and females was observed (Figure 4F, right panel). These findings indicate that the T-ALL-suppressive effect of the male antigen was dependent on restricted MHC class II molecules. To confirm that delayed Marilyn leukemogenesis in male mice was due to male antigen presentation, we next treated syngeneic immunodeficient females infused with Marilyn T-ALL with the DBY cognate peptide. Indeed, DBY administration significantly delayed T-ALL development in females (Figure 4F). Overall, these data indicate that strong agonistic HY-TCR signaling suppresses leukemogenesis.

## Discussion

In this report, we addressed the functional role of TCR signaling intensity in mouse T-ALL. We show that transgenic Marilyn HY-TCR tonic signaling not only had a synergistic effect with the ETV6::JAK2 fusion kinase, but also was sufficient to induce T-ALL on its own. Furthermore, our data show that the TCR expressed in Marilyn T-ALL cells responded to agonist stimuli, inducing apoptosis and leading to leukemogenesis suppression. Thus, these results support the notion that tonic TCR signaling can be leukemogenic, and strong signaling emanating from the same TCR is anti-leukemic.

Tonic signaling is generated by relatively weak TCR interactions with self-pMHC ligands and is important for controlling survival of naive T cells and allow their responsiveness to foreign antigens.^26^ Thymocytes expressing transgenic TCRs against foreign antigens (such as the male antigen for females) are subjected to self-pMHC-mediated tonic signaling during their maturation, namely at the positive selection stage. The notion that TCR signaling can be leukemogenic has been earlier described, and transgenic TCR mouse strains other than the Marilyn strain, were shown to lead to spontaneous T-ALL development with variable latency and penetrance. ^27–30^ It is thus likely that tonic signaling originates pro-survival or pro-proliferative genetic programs that can cooperate with stochastic oncogenic genetic alterations, such as *Notch1*-activating mutations and *Cdkn2a* deletions. *Cdkn2a* inactivation in particular was reported to facilitate the malignant transformation of immature T cells, ^31,32^ so it is possible that loss of *Cdkn2a* function may cooperate with TCR tonic signaling in DN thymocytes.

The finding that tonic TCR signaling can promote malignant transformation finds parallels in other systems. In fact, tonic B cell receptor signaling, defined as antigen-independent cell-autonomous signaling, has been shown to drive B cell lymphomagenesis ^33–35^ and to be therapeutically targetable.^36^ By the same token, tonic TCR signaling could be a targetable mechanism in human T-ALL, as suggested by the frequent activation of Lck kinase (a mediator of TCR signaling) in T-ALL and by the therapeutic potential of Lck inhibitors. ^6–9^ Lck kinase activation in T-ALL was attributed to constitutive pre-TCR signaling found in a subset of patients. ^6^ Since TCR and CD3 proteins are also frequently expressed in T-ALL, ^4^ further studies are warranted to determine the role of TCR signaling in human T-ALL pathogenesis.

Although Rag recombinase-mediated mutagenic events can contribute to leukemia development, ^37^ that was not the case for Marilyn T-ALL. In fact, T-ALL developed with higher penetrance and much faster in Marilyn;*Rag2*^−/−^ females than in Marilyn;*Rag2*^+/−^ females generated in the same animal facility. Since Rag2 deficiency arrests thymocyte development at the DN3 stage, these results suggest that ectopic TCR expression at the immature DN3 stage in the absence of more mature thymocytes has increased leukemogenic potential, in line with the finding that early expression of mature TCR in DN thymocytes promotes T-ALL development.^30^

Rag2-sufficient, Marilyn mouse cohorts housed in different animal facilities led to unexpected different rates of T-ALL incidence. Indeed, Marilyn T-ALL frequency was much higher in females housed at the UAlg/Faro facility than females housed at the i3S/Porto facility. Since several microorganisms (*Helicobacter* spp., *Pasteurella pneumotropica* and murine norovirus) were detected in the UAlg facility, but not in the i3S facility, increased exposure to pathogenic microbiota could have promoted a microenvironment favorable for transgenic TCR-induced leukemogenesis. This notion is supported by the finding of higher levels of LPS staining in lymphomas from UAlg facility mice than in lymphomas from i3S facility mice. Early research revealed that B-ALL in *Pax5* heterozygous mice was initiated only upon exposure to common pathogens. ^38^ Future studies should, therefore, determine whether the presence of microorganims promotes TCR-induced leukemogenesis and whether TCR interactions with MHC molecules presenting non-cognate foreign peptides generate pro-leukemogenic basal TCR signaling.

Although T-ALL in Marilyn females developed in the presence of self-pMHC, we found that MHC-dependent TCR signaling was not required for its maintenance. Indeed, secondary leukemia development was not affected by lack of MHC-bound self-peptides in recipient allogenic mice. These results contrast with the finding that continuous self-pMHC-mediated TCR signaling was necessary for lymphoma cell growth in a TCR-expressing reprogrammed T cell model. ^39^ Of note, expression of a transgenic TCR recognizing a peptide from the murine survivin protein was also shown to result in increased TCR signaling and to induce T-ALL. ^29^ In this case, the specific (self) antigen, survivin, was expressed constitutively, which contrasts with the absence of the HY-TCR cognate peptide (DBY) in Marilyn females. Our data together with previous reports demonstrate that the TCR signaling complex promotes T-ALL, and that this ability very much depends on the TCR affinity to antigen.

Finally, we showed that Marilyn T-ALL cells had a functional TCR and that *in vivo* TCR stimulation of transplanted female Marilyn T-ALL cells was associated with apoptosis and impaired Marilyn T-ALL leukemogenesis. These observations corroborate previous reports showing that mouse or human T-ALL underwent apoptosis upon activation of the TCR/CD3 signaling complex. ^19,28,40^ These findings suggest that, similarly to thymocytes undergoing thymic selection, TCR activation with low-affinity or high-affinity antigens will stimulate differentially the TCR signaling pathway and result in cell survival or death. ^41^ Future research should determine the TCR signaling regulators that dictate if a particular antigenic stimulus promotes or antagonizes T-ALL.

## Methods

### Mice

HY-TCR Marilyn transgenic mice (B6.Cg-Tg(TcraH-Y,TcrbH-Y)1Pas), on a CD45.1 background, were obtained from Jocelyne Demengeot (IGC, Oeiras). Marilyn mice were bred with EμSRα-ETV6::JAK2 (B6.Cg-Tg(Emu-ETV6/JAK2)71Ghy) transgenic mice ^13^, or *Rag2* (B6.129S6-Rag2^tm1Fwa^) knockout mice. The ETV6::JAK2 transgene was kept in hemizygosity. *Foxn1*^nu/nu^ (Nude) C57BL/6 and CBA mice were bred at the i3S facility. Mice were maintained at the UAlg/Faro and i3S/Porto barrier animal facilities, under 12 h light/dark cycles and with food and water *ad libitum*. HY-TCR Marilyn mice and ETV6::JAK2;Marilyn double transgenic were bred at the UALg animal facility, while the Marilyn;*Rag2*^+/−^ and Marily;*Rag2*^−/−^ were bred at the i3S animal facilty. Microorganism screening detected opportunistic pathogens (*Helicobacter* spp., *Pasteurella pneumotropica* and murine norovirus) in the UAlg, but not i3S, experimental rooms. All experimental procedures were approved by the i3S and CBMR/UAlg ethics committees and Portuguese authorities (*Direção-Geral de Agricultura e Veterinária*) and followed recommendations from the European Commission (Directive 2010/63/UE) and the local Portuguese authorities (*Decreto-Lei* n°113/2013). Both female and male mice were used for all experiments. Mice were monitored for signs of disease (e.g. dyspnea, lethargy, enlarged lymph nodes and enlarged abdomen) and killed by CO_2_ inhalation when reaching predefined experimental endpoints. Mice of different genotypes from the same litter were kept together in the same cages and monitoring for signs of disease was done blindly. Adult mice that were euthanized without leukemia were censored in Kaplan-Meier survival curves.

### Mice genotyping

The following primers were used for genotyping. HY-TCR Marilyn transgene: 5’-CGAGAGGAACCTGGGAGCTGT-3’ and 5’-TGCTGTCTGTACCACCAGAAATAC-3’; *Rag2*: 5’-TGTCCCTGCAGATGGTAACA-3’, 5’-CCTTTGTATGAGCAAGTAGC-3’, 5’-CTATTCGGCTATGACTGGG-3’ and 5’-AAGGCGATAGAAGGCGATG-3’; *Cdkn2a*: 5’-GTGATCCCTCTACTTTTTCTTCTGACTT-3’, 5’-CGGAACGCAAATATCGCAC-3’ and 5’-GAGACTAGTGAGACGTGCTACTTCCA-3’.

### *In vivo* experiments

For mouse leukemia transplantation assays, 0.5-2 x 10^6^ leukemic cells collected from diseased female Marilyn mice were intravenously injected in the tail vein of recipient 8-12-week-old C57BL/6, C57BL/6 Nude or CBA Nude mice of the indicated sex, and regularly monitored through peripheral blood detection of leukemic cells (CD45.1^+^CD45.2^+^TCR-Vβ6^+^). Female *Rag2*^−/−^ mice infused with Marilyn T-ALL cells were treated intraperitoneally once a week for two weeks with 0.01 mg of DBY peptide (NAGFNSNRANSSRSS; cat. no. AS-61046; Eurogentec).

### Flow cytometry

Single-cell suspensions were prepared from lymphoid organs using cell strainers, washed with FACS buffer (phosphate-buffered saline (PBS) with 3% fetal bovine serum (FBS) and 10 mM NaN_3_), and incubated for 30-45 min with fluorochrome-labeled antibodies in FACS buffer. The following Biolegend antibodies were used: CD25-fluoresceine isothiocyanate (FITC) (clone PC61), TCR β-phycoerithrin (PE) (clone H57-597), CD44-PE/Cyanine5 (clone IM7), CD90.2-PE-Cyanine7 (clone 30-H12), CD24-allophycocyanin (APC) (clone M1/69), CD4-APC/Cyanine7 (clone GK1.5), CD8 α-Pacific Blue (clone 53-6.7), Gr1-PE (clone RB6-8C5), CD127-PE/Cyanine5 (clone A7R34), B220-APC/Cyanine7 (clone RA3-6B2), CD24-Pacific Blue (clone M1/69), CD90.2-FITC (clone 30-H12), TCR-Vb6-PE (clone RR4-7), TCR-Vb5.1, 5.2-PE (clone MR9-4), CD5-APC (clone 53-7.3), CD69-FITC (clone H1.2F3), CD62L-APC (clone MEL-14), CD45.1-APC (clone A20) and CD45.2-Peridinin Chlorophyll Protein (PerCP)-Cyanine5.5 (clone 104). Immunostained cells were washed twice with FACS buffer and incubated in PBS with 10 mM NaN_3_. Cell viability was determined using the Zombie Aqua Fixable Viability Kit (Invitrogen). Samples were acquired using BD FACS Calibur, CANTO II or Accuri C6 and analyzed using FlowJo software.

### *Ex-vivo* T-ALL cell culture

Marilyn T-ALL cells were cultured ex-vivo for 16 h in RPMI medium supplemented with 10% FBS, 1% penicillin-streptomycin and 50 μM of 2-mercaptoethanol, all from Gibco. For *ex vivo* stimulation of Marilyn T-ALL cells, plates were coated with 10 μg/ml anti-CD3 (145-2C11, Biolegend) for 2 h at 37 °C, and then plates were washed with PBS. Ten ng/ml of PMA and 250 ng/ml Ionomycin (both from Sigma-Aldrich) treatments were performed for 18 hours.

### *Notch1* mutation detection

Genomic DNA from mouse leukemia samples was isolated using the GeneJET Genomic DNA purification kit (Thermo Fisher Scientific), following the manufacturer’s instructions. DNA was used for PCR amplification of two segments of *Notch1* exon 34: primer pair 5’-GCTCCCTCATGTACCTCCTG-3’ and 5’-TAGTGGCCCCATCATGCTAT-3’, generating a predicted amplicon of 904 bp, and primer pair 5’-ATAGCATGATGGGGCCACTA-3’ and 5’-CTTCACCCTGACCAGGAAAA-3’, generating a predicted amplicon of 893 bp. PCR products were Sanger sequenced using the same primers at i3S Genomics (Porto, Portugal) or CCMAR *Serviços de Biologia Molecular* (Faro, Portugal).

### Low-coverage whole genome sequencing

Genomic DNA was used for copy number analysis performed as described previously^42^ using a HiSeq 4000 (Illumina) sequencer in a single-read 50-cycle run mode. For copy number analysis, sequencing data was aligned with the mouse genome (mm10) as reference and further analyzed using QDNAseq (RRID:SCR_003174) package (version 1.12) in R software. A bin size of 15 kb was used.

### Histology and immunohistochemistry

For histological and immunohistochemistry analysis, formalin-fixed paraffin-embedded tissues were sectioned with 4 μ m thickness. Hematoxylin and eosin staining was performed using standard procedures. For immunohistochemistry, heat-mediated antigen retrieval was performed for 35-40 min with citrate-based antigen retrieving solution (cat. no. H-3300, Vector Laboratories), for cleaved-caspase 3, Ki67 and LPS immunodetection, or 10 μM EDTA, pH 8.0, and 0.05% Tween 20 in PBS, for B220 or CD3 immunodetection. Endogenous peroxidase was inactivated with 3% H_2_O_2_ in methanol, and nonspecific antibody binding was blocked using Ultravision Protein-block (Thermo Fisher Scientific). Sections were incubated overnight with rat anti-B220 (1:200, clone RA3-6B2, Biolegend), anti-CD3 (1:200, ab5690, Abcam), anti-cleaved-caspase 3 (1:200, 9661, Cell Signaling), Ki67 (1:500, ab15580, Abcam) or LPS (1:100, WN1 222-5, Hycult Biotech) at 4 ° C. For B220 detection, slides were incubated sequentially with avidin/biotin blocking system (Biolegend), goat anti-rabbit secondary antibody (Biovision), and streptavidin-horseradish peroxidase (HRP) (Enzo Biochem). For CD3, Ki67 and LPS detection, HRP-conjugated rabbit/mouse secondary antibody (Agilent) was used. 3,3′-Diaminobenzidine chromogen (Agilent) was used as detection reagent. For quantification of percentage of cleaved caspase 3-positive cells, five images were taken from different areas of each tissue section.

### Statistics

Statistical analysis was performed with GraphPad Prism 6.0 software (RRID:SCR_002798). Log-rank test was used to compare survival of different groups. Unpaired student’s t-test was used for comparisons between two groups. Welch’s correction was used when variances were different. Mann-Whitney test was used to compare white blood cell (WBC) count. Detection of CD45.1^+^ T-ALL cells in the peripheral blood of males and females was compared using two-way ANOVA. Sample numbers are indicated in figure legends. *P*<0.05 was considered statistically significant.

## Data Sharing Statement

The data that support the findings of this study are available from the corresponding author, upon reasonable request. Raw unaligned sequencing reads (fastq-format) that support the findings of this study have been deposited in the SRA.

## Acknowledgments

We thank Jocelyne Demengeot (IGC, Oeiras) and Nuno L. Alves (i3S, Porto) for providing Marylin transgenic mice. We thank members of i3S Intercellular Communication and Cancer group for fruitful discussions about this work. The authors acknowledge the support the Animal Facility, Translational Cytometry and Histology, Electron Microscopy (member of the Portuguese Platform of Bioimaging; PPBI-POCI-01-0145-FEDER-022122) and Genomics i3S Scientific Platforms. This work was supported by a fellowship (FAZ Ciencia prize) from Fundação AstraZeneca (Lisbon, Portugal). This work was supported by European Regional Development Fund (ERDF), through COMPETE 2020 - Operational Program for Competitiveness and Internationalisation, Portugal2020, and *Fundação para a Ciência e a Tecnologia* (FCT; POCI-01-0145-FEDER-007274; PTDC/MED-ONC/32592/2017), and by ERDF, through the Norte Portugal Regional Program (NORTE2020), Portugal2020 (NORTE-01-0145-FEDER-000029) and HEALTH-UNORTE (NORTE-01-0145-000039). T.A.C. was recipient of FCT (PD/BD/114129/2015) and *Liga Portuguesa Contra o Cancro – Núcleo Regional do Norte* fellowships. J.L.P. (SFRH/BD/147979/2019) and M.B. (2021.06024.BD) were recipients of FCT fellowships.

## Contribution

T.A.C. designed, performed and analyzed experiments, made the figures and wrote the paper; I.P.-L., J.L.P., and M.B. performed experiments; N.R.S. designed the study, performed experiments, made the figures and wrote the paper.

