## Supplemental Figures and Tables for "Transgenic αβ TCR tonic signaling is leukemogenic while strong stimulation is leukemia-suppressive" (from another version of this preprint: version 1, 2024-01-22)

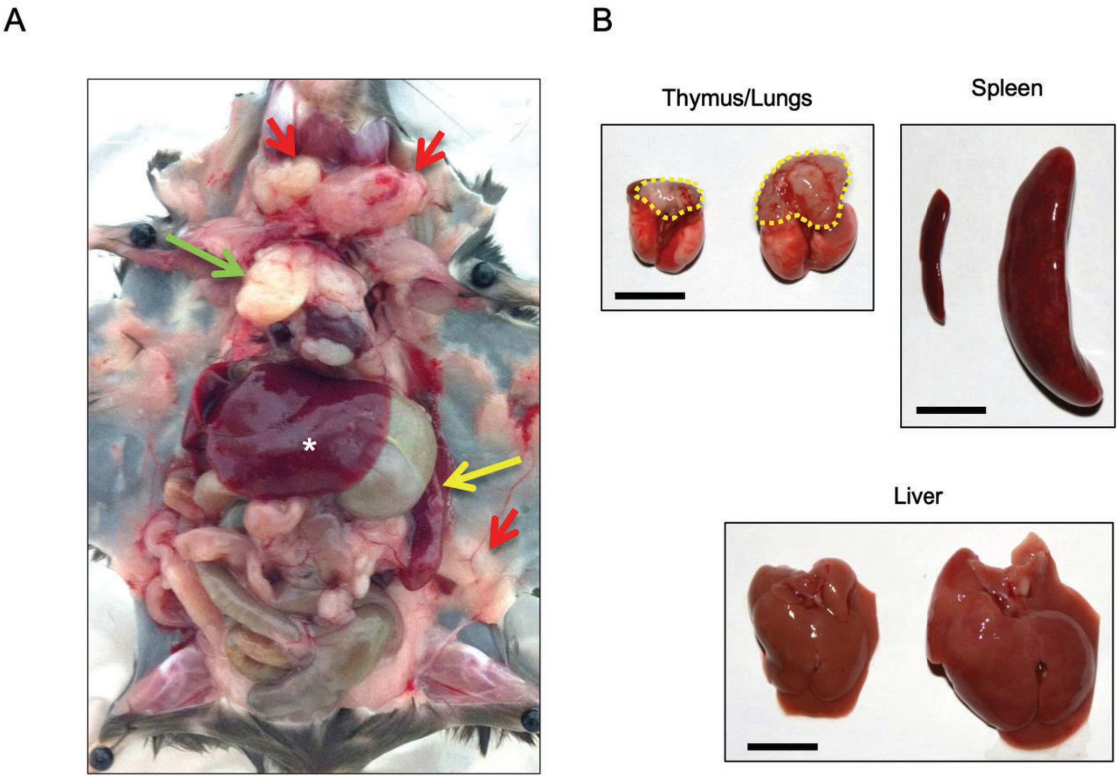


**Supplementary Figure 1. Marilyn female mice developed hematological tumors.** (A) Representative female Marilyn mouse (no. 34) presenting enlarged thymus (green arrow), LNs (red arrows) and spleen (yellow arrow). (B) Enlarged thymus, spleen and liver from a female Marilyn mouse (no. 3), compared to wild-type mouse organs. Yellow dotted line highlights the thymus. Scale bar: 1 cm.

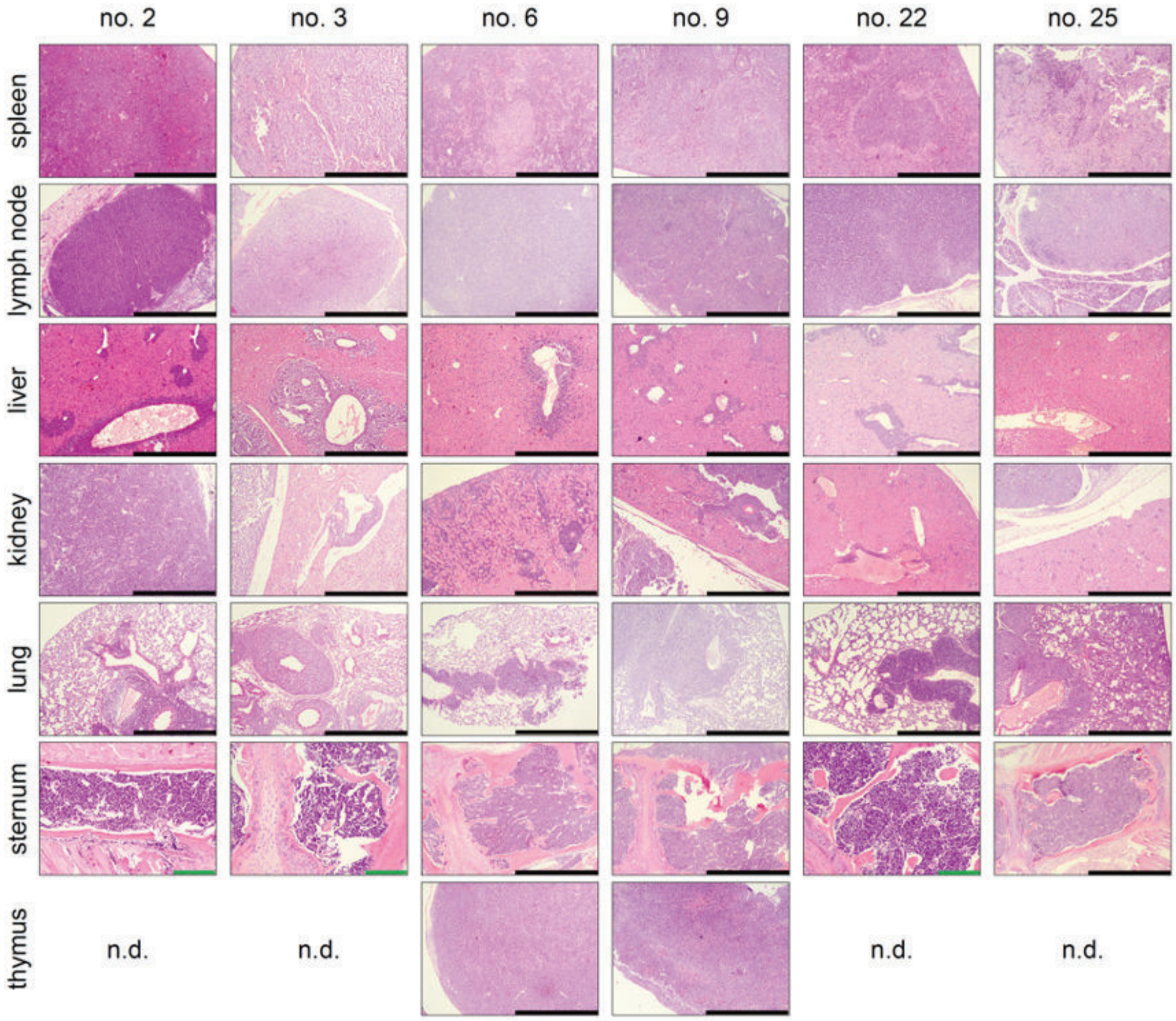


**Supplementary Figure 2. Histological analysis of Marilyn female mice with leukemia/lymphoma.** Hematoxylin and eosin staining of sections of the indicated organs of diseased Marilyn female mice. Black scale bars: 500 μm; green scale bars: 200 μm. n.d., not determined.

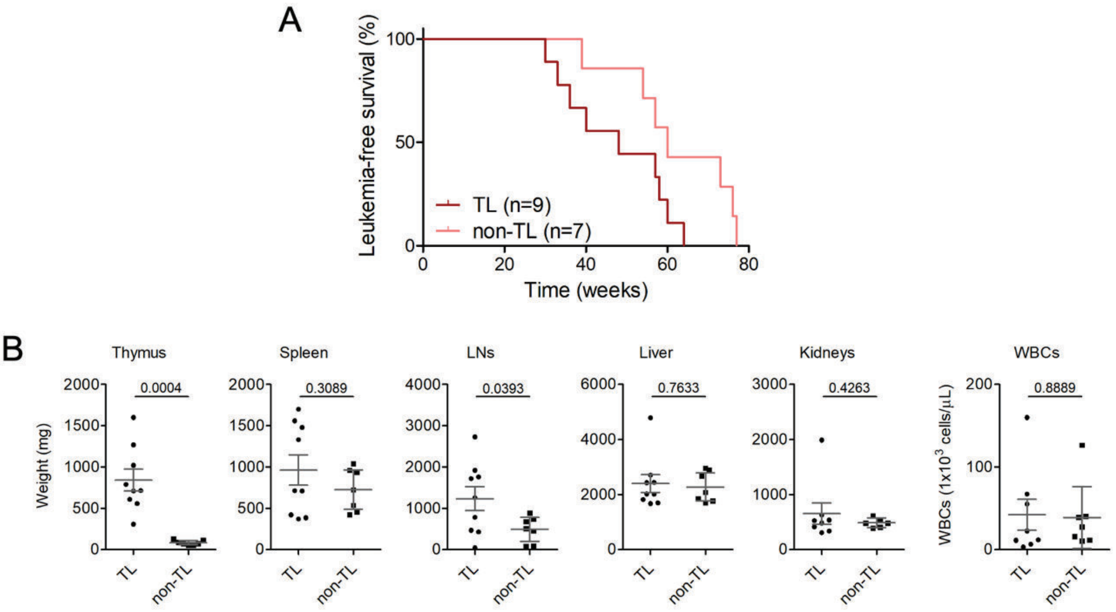


**Supplementary Figure 3. Marilyn female mice with thymic lymphoma lived shorter and presented more severe lymphadenopathy than remaining diseased mice.** (A) Kaplan-Meier survival curves for female with thymic lymphoma (TL) or developing lymphoma in organs other than thymus (non-TL). *P*=0.0509, log-rank test. (B) Organ weights and white blood cell counts (WBCs) of mice developing thymic lymphoma versus non-thymic lymphomas. *P* values were determined by unpaired *t* tests, with Welch's correction when variances were significantly different. Mean and SEM are shown.

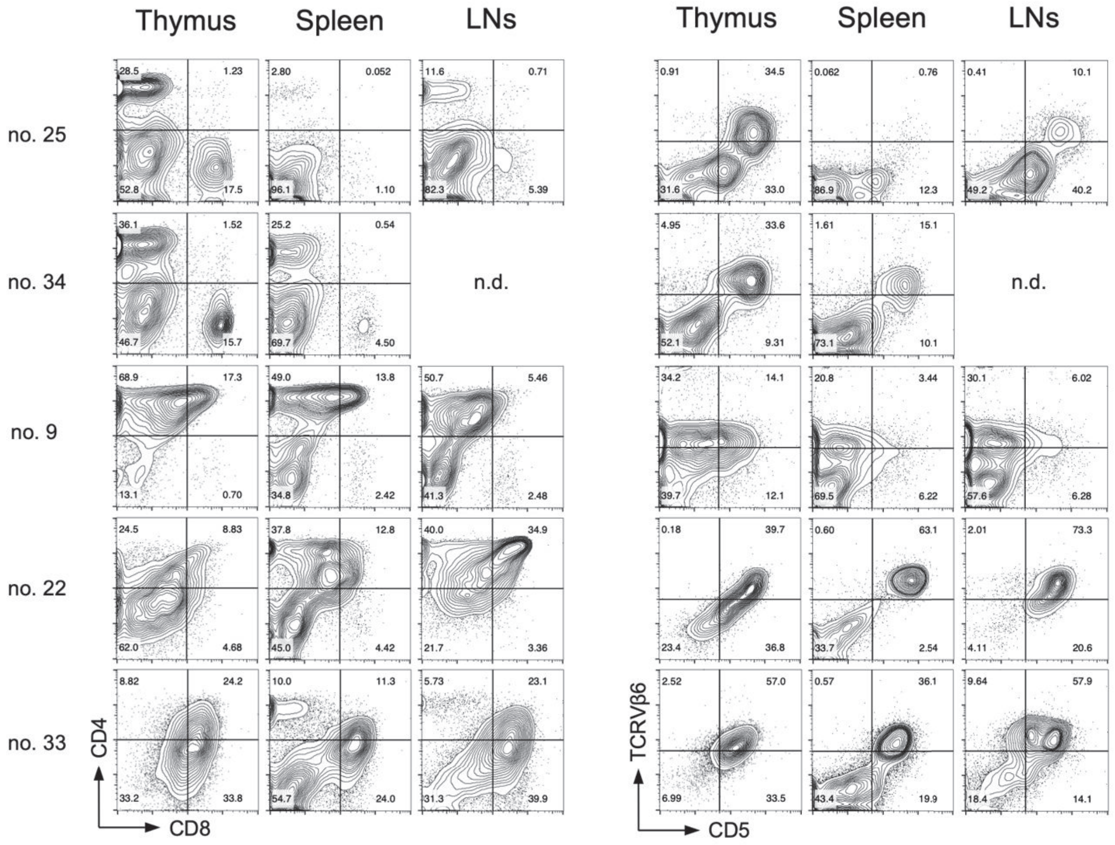


**Supplementary Figure 4. The immunophenotype of pathological HY-TCR T cells is similar among different lymphoid organs of the same individual.** Flow cytometry detection of CD4, CD8, TCR-Vβ6 and CD5 surface expression in thymus, spleen and LNs from the indicated diseased Marilyn females. n.d., not determined.

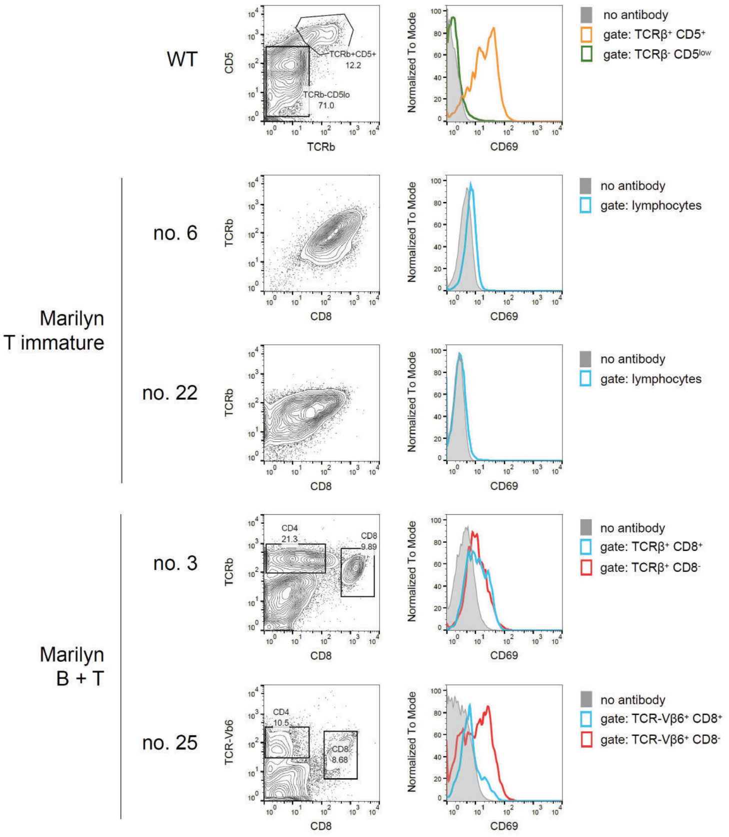


**Supplementary Figure 5. CD69 expression in pathological HY-TCR T cells.** Flow cytometry analysis of TCRβ, TCR-Vβ6, CD8 and CD69 expression for wild-type (WT) thymocytes and thymic cells from 4 cases of diseased Marilyn females ascribed to the immature T or B+T disease subtypes. Right panels show CD69 surface expression in positively selected ($TCR\beta^{+}CD5^{+}$) WT thymocytes, all leukemic cells (cases nos. 6 and 22) or TCR-expressing cells (cases nos. 3 and 25).

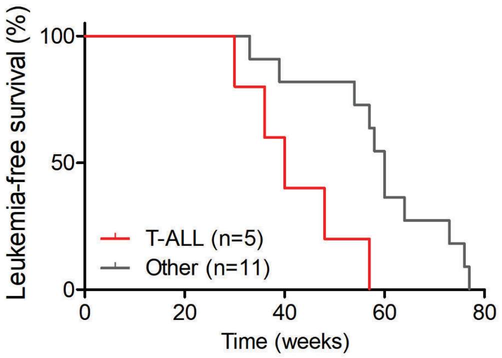


**Supplementary Figure 6. Marilyn female mice with T-ALL showed earlier disease onset.** Kaplan-Meier survival curves for females diagnosed with either T-ALL or other hematological disease. $P$=0.0076, log-rank test.

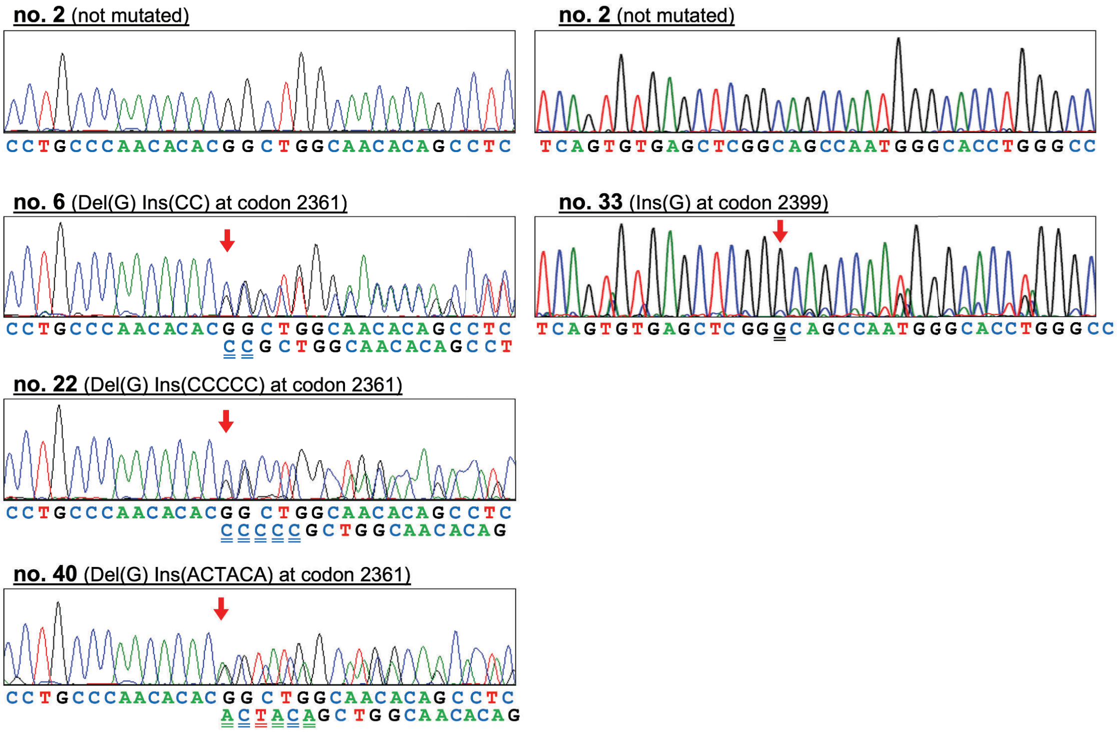


**Supplementary Figure 7. *Notch1* mutations were detected in immature type Marilyn leukemic cells.** Sanger sequencing chromatograms of *Notch1* exon 34 (encoding the PEST domain) fragments for Marilyn T-ALL cells. Thymic leukemic cells from mice nos. 6, 22, and 40 carried heterozygous frameshift mutations in codon 2361, as revealed by the presence of overlapping peaks, representing two parallel sequences. Thymic leukemic cells from mouse no. 33 carried a homozygous frameshift mutation in codon 2399, while those from mouse no. 2 had no PEST domain mutation. Mutation localization is indicated by a red arrow.

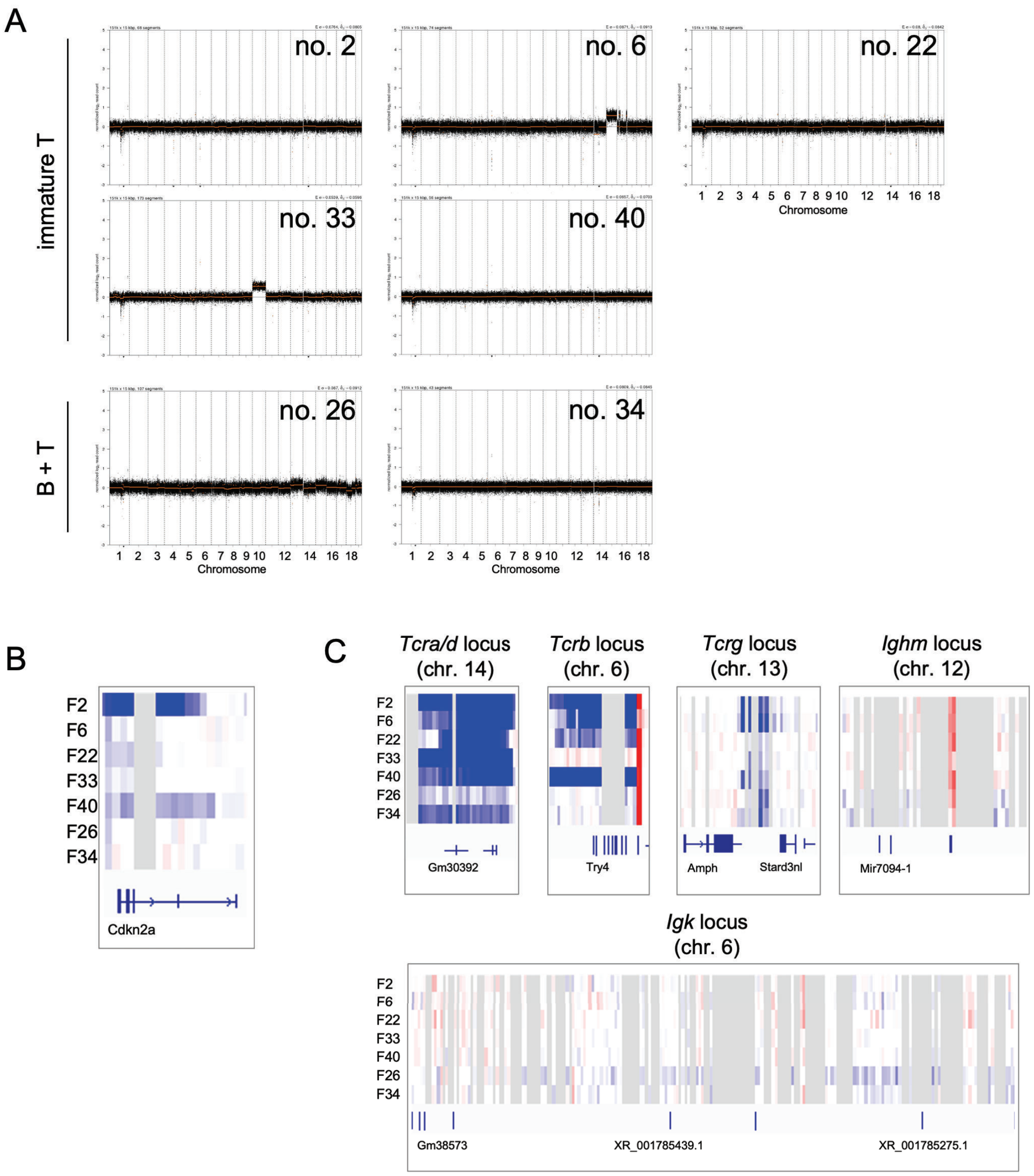


**Supplementary Figure 8. DNA copy number alterations detected in Marilyn T-ALL cells.** (A) DNA copy number alterations in leukemic cells from representative female Marilyn mice assessed by low coverage whole-genome sequencing. Each graph represents DNA copy number distribution for each case, with signals above and below zero representing respectively DNA gains and DNA losses. (B) DNA copy number variation analysis shows deletions in the *Cdkn2a* locus in two cases (nos. 2 and 40) of Marilyn T-ALL. (C) DNA copy number variation analysis for *Tcra/d*, *Tcrb*, *Tcrg*, *Ighm* and *Igk* loci in cells from diseased Marilyn mice. In B and C blue represents copy number loss and red represents copy number gain relative to reference mouse genome (mm10).

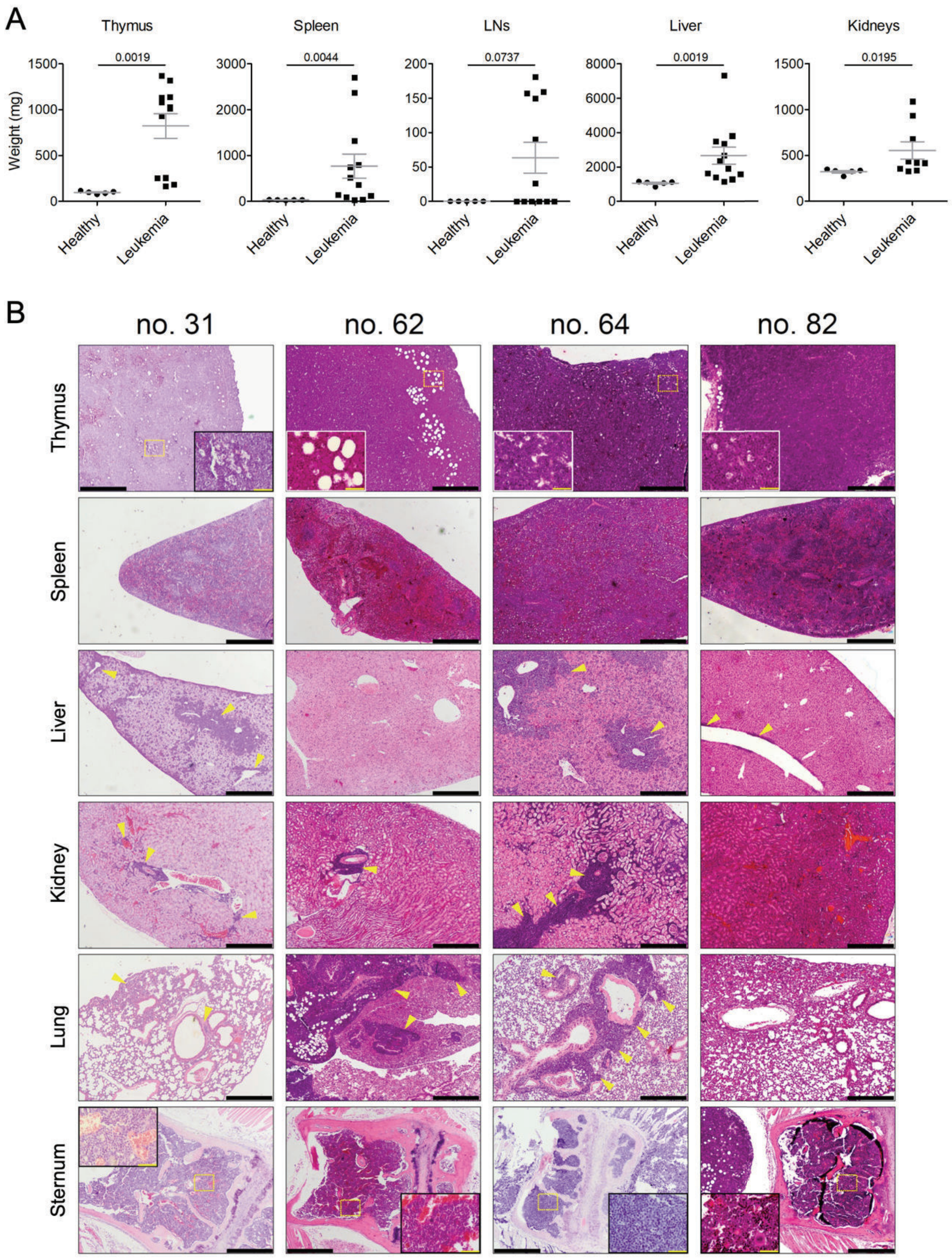


**Supplementary Figure 9. Transgenic HY-TCR expression in *Rag2*$^{-/-}$ female mice leads to development of aggressive T-ALL disseminated to several organs.** (A) Organ weights of Marilyn;*Rag2*$^{-/-}$ females with leukemia/lymphoma. Healthy females were used as control. *P* values determined by unpaired *t* tests with Welch's correction. (B) Hematoxylin and eosin staining of sections of the indicated organs of Marilyn;*Rag2*$^{-/-}$ females with T-ALL. Yellow arrowheads indicate tissue areas infiltrated with leukemic cells. Black scale bars, 500 μm; yellow scale bars, 200 μm.

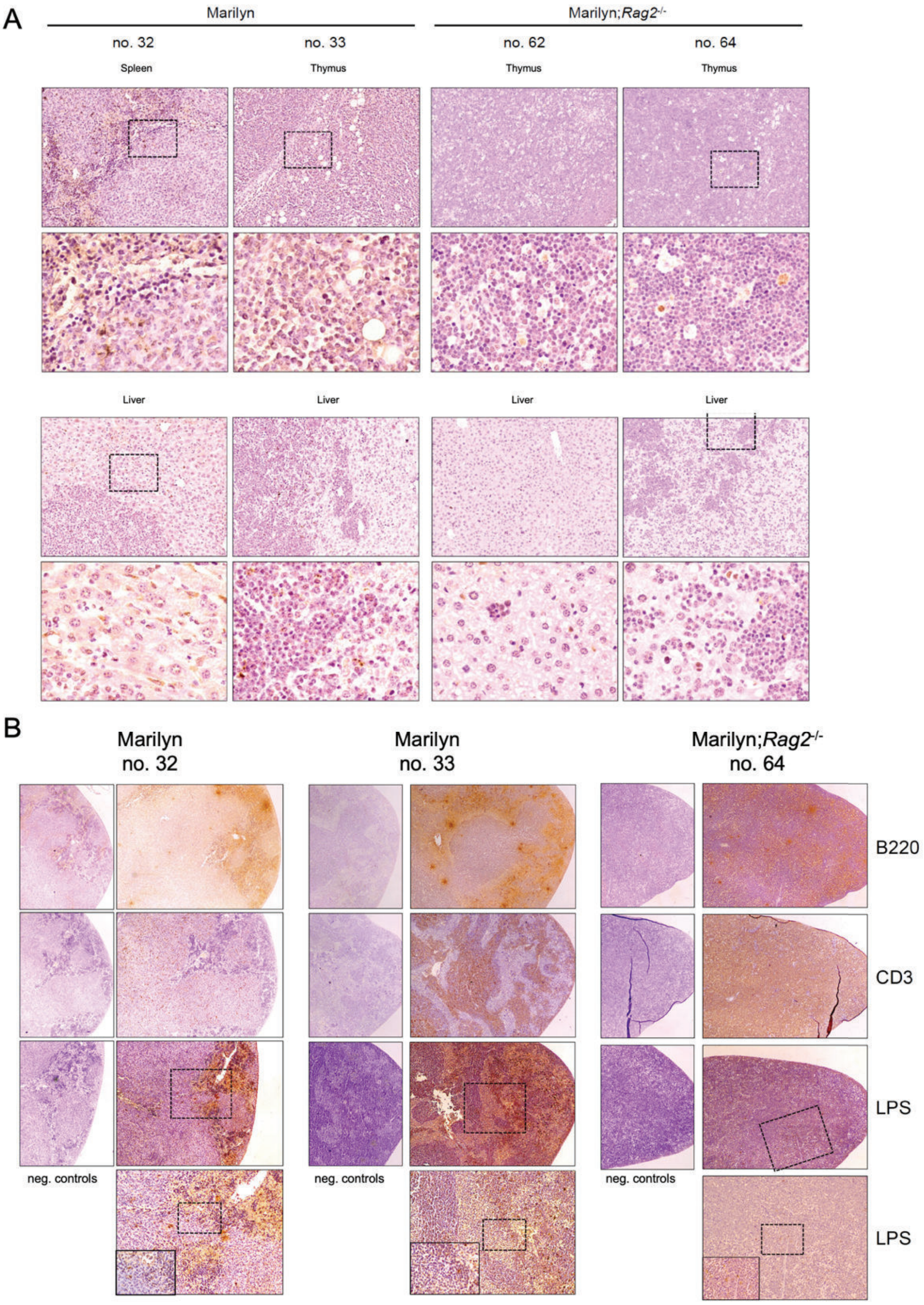


**Supplementary Figure 10. Lipopolysacharide immunostaining in organs from diseased Marilyn leukemic females.** (A) Anti-lipopolysacharide (LPS) immunohistochemistry for livers and thymus (or spleen for no. 32) of representative Marilyn and Marilyn;*Rag2*$^{-/-}$ mice, housed in the UAlg/Faro and i3S/Porto facilities respectively. Top panels: 100x magnification; bottom panels: 400x magnification. (B) LPS, CD3 and B220 immunohistochemical detection in splenic lymphoma sections of representative Marilyn or Marilyn;*Rag2*$^{-/-}$ females. Staining with secondary reagent without primary antibody was performed as negative control (left panels for each mouse sample). Top 3 rows: 40x magnification; bottom row: 100x magnification; inset: 400x magnification.

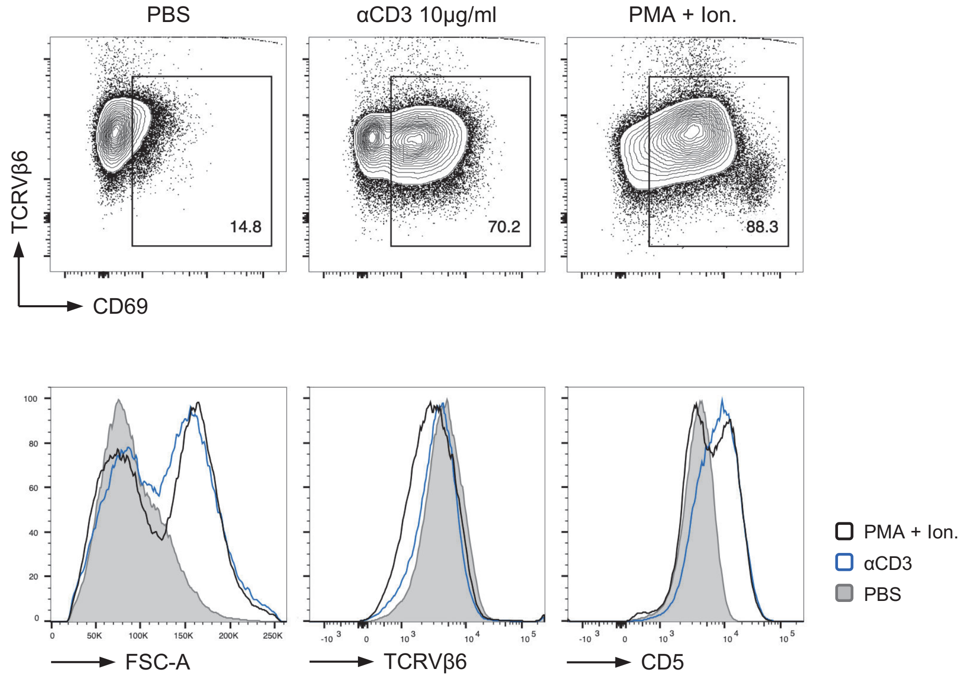


**Supplementary Figure 11. Marilyn T-ALL cells are responsive to TCR signaling activation.** Flow cytometry detection of TCR-Vβ6, CD5 and CD69 expression in Marilyn T-ALL cells stimulated *ex vivo* with 10 μg/mL plate-bound anti-CD3 (αCD3) or 10 ng/mL PMA and 250 ng/mL ionomycin (Ion.) for 16 hours. Bottom histograms show forward side scatter (to gauge cell size) and expression of TCR-Vβ6 and CD5 in stimulated CD8+ Marilyn T-ALL cells. Data are representative of 3 independent experiments performed.

**Supplementary Table 1.** Macroscopic, immunophenotypic and molecular features of diseased Marilyn female mice.

| Mouse no. | Age (wks) | Thymus (mg) | Spleen (mg) | LN (mg) | WBC (cells/µl) | BM infiltration | Predominant Immunophenotype | Notch1 exon 34 | Copy number changes | Transplantability |
|---|---|---|---|---|---|---|---|---|---|---|
| Thymic lymphoma (immature T phenotype) | | | | | | | | | | |
| **2** | 40 | 560 | 1,330 | 1,260 | - | - | T immature | WT | *Cdkn2a* loss | 6/6 |
| **6** | 57 | 1,270 | 1,560 | 1,920 | 6,500 | 95% | T immature | MUT | +Chr.15; +Chr.16 | 4/4 |
| **22** | 36 | 715 | 710 | 2,730 | 55,000 | 87% | T immature | MUT | None | 4/4 |
| **33** | 48 | 1,020 | 385 | 430 | 22,500 | 8% | T immature | MUT | +Chr.10 | 5/6 |
| **40** | 31 | 1,600 | 375 | 40 | 11,900 | 4% | T immature | MUT | *Cdkn2a* loss | 4/5 |
| Thymic lymphoma (mature T phenotype) | | | | | | | | | | |
| **3** | 60 | 610 | 170 | 1,720 | 11,300 | 6% | B + T mature | WT | - | 0/2 |
| **9** | 64 | 310 | 715 | 1,760 | 160,000 | 8% | T mature | - | - | 0/2 |
| **29** | 33 | 710 | 1,480 | 470 | 67,200 | 75% | T mature | - | - | 0/4 |
| **34** | 58 | 790 | 420 | 785 | 3,000 | 6% | B + T mature | WT | None | - |
| Non-thymic lymphoma | | | | | | | | | | |
| **11** | 73 | 115 | 420 | 675 | 39,300 | 73% | B + T mature | - | - | - |
| **12** | 76 | 50 | 725 | 90 | 27,600 | 7% | B + T mature | - | - | - |
| **25** | 39 | 130 | 960 | 505 | 126,400 | 4% | B + T mature | - | - | 0/2 |
| **26** | 60 | 50 | 535 | 455 | 11,500 | 8% | B + T mature | WT | +Chr.13,15; -Chr.18 | - |
| **31** | 54 | 70 | 460 | 75 | 40,000 | 3% | B + T mature | - | - | - |
| **32** | 77 | 75 | 940 | 740 | 10,400 | 5% | B + T mature | WT | - | - |
| **38** | 57 | 90 | 1,040 | 890 | 15,800 | 5% | B + T mature | WT | - | - |

-, not determined; MUT, mutation; WT, wild-type sequence

**Supplementary Table 2.** Macroscopic, immunophenotypic and molecular features of diseased Marilyn;*Rag2*$^{-/-}$ female mice.

| Mouse no. | Age (wks) | Thymus (mg) | Spleen (mg) | LN (mg) | WBC (cells/µl) | BM infiltration | Immunophenotype | *Notch1* exon 34 | *Notch1* mutated codon | Transplantability |
|---|---|---|---|---|---|---|---|---|---|---|
| **14** | 42 | 1130 | 138 | 0 | - | 11% | DN+DP, Vβ6$^{+}$, CD5$^{+}$, CD24$^{+}$, CD44$^{+}$ | MUT | 2361, del G, ins CC | 0/2 |
| **26** | 39 | 930 | 355 | 90 | 37,800 | 52% | DN+DP, Vβ6$^{+}$, CD5$^{low}$, CD24$^{+}$ | MUT | 2418, ins TAGCTTTA | - |
| **28** | 35 | 254 | 1320 | 159 | 26,600 | 91% | DP+CD8SP, Vβ6$^{+}$, CD5$^{+}$, CD24$^{+}$ | MUT | 2399, ins AGGGTA | - |
| **31** | 34 | 1370 | 120 | 0 | 36,400 | - | - | WT | - | 1/1 |
| **33** | 46 | 164 | 740 | 0 | 50,800 | - | DP+CD4SP, Vβ6$^{+}$, CD5$^{+}$, CD24$^{+}$, CD69$^{+}$ | WT | - | - |
| **58** | 60 | 1030 | 800 | 0 | - | - | - | MUT | 2490, del CT | - |
| **60** | 67 | 183 | 2700 | 150 | 2,350 | 18% | DN+DP, Vβ6$^{+}$, CD5$^{low}$, CD24$^{+}$ | MUT | 2490, del CT | - |
| **62** | 50 | 1140 | 32 | 26 | 4,700 | - | - | MUT | 2361, del G, ins CTC | 4/5 |
| **64** | 46 | 1080 | 516 | 157 | 356,800 | - | - | WT | - | 5/5 |
| **80** | 64 | 255 | 2370 | 181 | 131,200 | - | - | - | - | 5/5 |
| **81** | 69 | 1320 | 36 | 0 | 1,700 | - | DN+CD8SP, Vβ6$^{+}$, CD5$^{+}$ | - | - | - |
| **82** | 38 | 1014 | 90 | 0 | 7,800 | - | - | WT | - | - |

-, not determined; DN, CD4/CD8 double-negative; DP, CD4/CD8 double-positive; MUT, mutation; WT, wild-type sequence; del, deletion; ins, insertion
